# Performance and limitations of out-of-distribution detection for insect DNA (meta)barcoding

**DOI:** 10.1101/2025.10.19.683333

**Authors:** Tomochika Fujisawa, Takashi Imai

**Author notes:** Corresponding author Tomochika Fujisawa. Data availability: A full list of BOLD accession numbers and sequence alignments used in this study are available at http://doi.org/10.6084/m9.figshare.c.8021959, The code to reproduce this study is available at https://github.com/tfujisawa/barcoding_cnn. Conflict of interest: The authors declare no conflict of interest.

## Abstract

Successful applications of DNA barcoding/metabarcoding rely on the accurate taxonomic identification of sequence fragments. When biological surveys with DNA (meta)barcoding target underexplored biological communities, sequence-based identification is often conducted using incomplete databases that do not fully cover the regional species pool. Consequently, specimens to be identified may include species not present in reference databases. Such unknown or “out-of-distribution” samples can cause misidentification if left undetected. A similarity cutoff is commonly used to detect out-of-distribution samples before taxonomic assignment, but its effectiveness has not been carefully studied. In this study, we evaluated the performance of out-of-distribution detection for DNA barcoding with genetic distance and deep learning metrics. Using extensively sampled datasets of multiple insect taxa, we measured the performance of identification and out-of-distribution detection under conditions in which genetic variations in species were sufficiently sampled. Although identification with DNA barcoding is a highly accurate process, even with short noisy fragments, out-of-distribution detection was more susceptible to a reduction in performance due to sequence noise and a lack of diagnosable characters. Our results provide guidelines for designing unknown-proof identification procedures by determining factors affecting out-of-distribution detection performance.

## INTRODUCTION

The reliable identification of specimens to known taxonomic groups is the foundation of biological studies. Without accurate identification, subsequent practices, including conservation, biosecurity, and ecological monitoring may not be conducted reliably. Despite its importance, taxonomic expertise is a scarce resource, and the identification of specimens is a major bottleneck in large-scale ecological surveys (Van Klink et al. 2024). Driven by the general lack of taxonomic expertise and the need for rapid and broad characterization of threatened biodiversity, replacing or complementing human identification with computational methods has attracted attention in recent decades (MacLeod et al. 2010; Gaston & O’Neil 2004). In particular, methods based on DNA sequences have been considered promising because they can enable high-resolution, species-level identification that is accessible to non-experts. DNA barcoding (Hebert et al. 2003), which is the process of identification based on standardized short DNA fragments (e.g., CO1 fragments for animal identification), is the most successful project of such attempts. Currently, the Barcode of Life Data system has 16 million registered sequences, and identification using barcoding markers is routinely performed. The recent introduction of high-throughput sequencing technologies has broadened the scope of DNA barcoding applications. A notable example is DNA metabarcoding (Taberlet et al. 2012), parallel sequencing and identification of barcoding markers, either from a bulk sample of organisms or environmental DNA. Metabarcoding has significantly expanded the scale of biological monitoring by increasing throughput (Srivathsan et al. 2019) and has widened research targets to previously neglected communities, such as meiofauna or soil arthropods (Macher et al. 2024; Depheide et al. 2019).

Although promising, modern applications of DNA metabarcoding are practiced under conditions that are substantially different from those for which DNA barcoding was initially designed, posing new methodological challenges. For example, identification is conducted using fragments shorter than the original barcoding markers (originally up to 1000 bp, but now shorter than 400bp) (Leese et al. 2021; Miya et al. 2015; Leray et al. 2013) because of the limitations associated with efficient PCR amplification and high-throughput sequencing.

Taxonomic identities are often recovered under noisy conditions involving more sequencing errors and artifacts. In addition, metabarcoding surveys target largely unexplored biota whose members are undescribed and there is a lack of representative sequences in reference databases. Hence, the re-optimization of identification procedures has ensued since the introduction of high-throughput technologies, including molecular protocols and bioinformatic pipelines (Creedy et al. 2021; Alberdi et al. 2017).

One aspect of metabarcoding applications requiring close attention is the effect of samples of the class not present in the reference database. Samples of classes that are not present in the reference are called by different names in different application fields, reflecting the nature of such samples: unknown, novelty, anomaly, outliers and out-of-distribution. We use out-of-distribution (hereafter, OOD) samples in this study because it is a general term that sufficiently encompasses our task. Also, in the context of sequence-based taxonomic identification, absence from a reference dataset does not immediately define the exact nature of a sample such as “novelty”.

It has been shown that the current sequence databases do not fully represent the diversity of life, and this trend is especially prominent for highly diverse groups such as arthropods. According to previous studies, less than 20% of described invertebrate species have sequence records in public repositories (Keck et al. 2023), and only approximately 20% of terrestrial arthropod species have been formally described (Stork 2018). The use of underrepresented databases for metabarcoding surveys inevitably results in OOD samples. Indeed, large-scale metabarcoding applications have reported many unknown species, even from among supposedly well-studied fauna (Buchner et al. 2024). Because current reference databases are incomplete and undetected OOD samples certainly cause misidentification, molecular identification methods under metabarcoding projects require the appropriate handling of OOD samples. In short, any identification algorithm should be able to say, “I do not know.”

The treatment of OOD samples has been an important issue for practical applications involving DNA barcoding because encountering them is the norm rather than an exception in most conditions. Protocols for detecting unknowns with distance thresholds have been considered even in the very early stages of DNA barcoding studies (Meier et al. 2006). Empirical thresholds with sequence similarity are still commonly used to remove putative unknown samples or retain them for higher-class assignments (e.g. a 97% similarity threshold). This approach also includes thresholds using reliability scores instead of distance, such as bootstrap uncertainty scores (Murali et al. 2018; Porter et al. 2014). More recently, methods that explicitly model the encounters of unknown or new species have been introduced. Methods such as PROTAX and BayesANT (Zito et al. 2023; Somervuo et al. 2016) explicitly model the probability of finding unknowns under species sampling models and infer the sample’s posterior probability of being an unknown species.

Nevertheless, apart from these studies, performance evaluations of OOD detection procedures have not often been conducted systematically. Performance evaluation often does not enable us to distinguish between the failure of OOD detection and misidentification of in-distribution samples, even if these two types of errors represent failures at different steps in the identification process. Critical issues, such as the methods that are favorable for use under certain conditions, have not been clearly addressed, and the major determinants of detection performance are unknown.

In this study, we evaluated molecular taxonomic assignment methods for DNA barcoding, with a specific focus on the effects of incomplete databases and the presence of OOD samples. We used insect DNA barcoding data to conduct the performance evaluation. Insects are major targets of DNA barcoding projects because of their extreme diversity and the difficulties associated with manual identification. Recent large-scale inventorying efforts (Roslin et al. 2022; Hebert et al. 2016) have enabled performance testing under ideal conditions, where a sufficient number of samples are available to characterize species genetic variations across clades. The large sample size also enables the training of parameter-rich machine learning models, including deep learning models. Recently, deep learning has been successfully applied to various biological sequence analyses, including taxonomic classification (Romeijn et al. 2024; Ziemski et al. 2021; Busia et al. 2018), but its performance in incomplete databases is unexplored.

We tested the performance of conventional and deep learning algorithms for taxonomic assignment and OOD detection using extensively sampled insect taxa. We explored the performance limits of identification and OOD detection methods by focusing on insect taxa with sufficient within- and between-species sampling. We showed that both conventional and deep learning identification methods are highly accurate for taxonomic assignment and robust for short and noisy sequences, whereas OOD detection is more prone to performance reduction due to noise and limited availability of information in short fragments.

## MATERIALS AND METHODS

### Data acquisition

We tested the performance of the classification and OOD detection models using datasets downloaded from the Barcode of Life Data (BOLD) System database (Ratnasingham & Hebert 2007). We first conditioned database entries by geographic regions where comprehensive inventorying of regional insect fauna was underway (mainly North American and EU countries), and then selected insect genera with sufficient sample size, taxonomic coverage, and geographic extent. Genera were selected as targets when at least 15 species were represented by 15 or more individuals (Fifteen individuals within species is a sample size large enough for correct estimation of within-species genetic variations upon training) (Zhang et al. 2010; Matz & Nielsen 2005). We applied this criterion to four major insect groups, i.e., Hymenoptera (bees and wasps), Diptera (flies), Lepidoptera (moths and butterflies), and Coleoptera (beetles), as these groups had the densest and broadest samples in the BOLD database. Then, twenty candidate genera were randomly selected. Classification models were trained to conduct species-level identification within the genus. A full list of BOLD accession numbers and sequence alignments are available in the Supplementary Data.

### Data preparation

The downloaded CO1 sequences of the 20 genera were filtered according to length and sequenced regions. Only fragments with length > 400 bp and < 1000 bp and fragments with the “5BP” DNA barcoding region were retained. Sequences with missing bases in the latter half of the 5BP region were discarded because they did not contain the short barcoding regions used for performance evaluation. Samples identified only at the genus level were removed, but samples with unconventional labels, such as “sp. 10” or “sp. DNAS-***” were included in OOD datasets because these names were consistently applied to multiple samples and likely represented true unknown species. To reduce the adverse effects of overrepresented species, species with >125 samples were randomly resampled to reduce the number of samples to 125, which was sufficient to characterize the genetic and haplotype diversity of the focal species (Zhang et al. 2010).

The filtered sequences were then aligned using MAFFT (v.7.453, Katoh et al. 2013) with default parameters. Aligned sequences were split into in-distribution (ID) and OOD samples based on the number of individuals in the species (based on the logic that rare species are more likely to be OOD). Species with ≥ 15 samples were assigned to ID and the other species were assigned to OOD. Classification models were first trained to classify ID samples into their taxonomic groups and were subsequently exposed to OOD samples to test whether the models could correctly detect them.

We compiled multiple datasets covering various parameters, including the total number of samples, fragment lengths, and sequencing errors. We prepared two short alignments by selecting the 350-650 region (300 bp) and 350-500 (150 bp) within full alignments. These 300 bp and 150 bp regions largely overlap with the short barcoding regions proposed by Lelay et al. (2013) and Leese et al. (2021), respectively. We also randomly halved the number of samples to create smaller datasets in which at least five samples per species were retained in the training process. We named these halved datasets “*Small*” and original datasets “*Sufficient*.” To simulate sequencing errors, bases were randomly swapped with a noise rate of 0.02, where randomly selected 2% of bases in a fragment were replaced with one of the alternative bases with an equal probability of 1/3. Errors were introduced only into test datasets because only clean reference databases were available for model training in realistic applications. The final datasets covered two database size categories {“Sufficient,” “Small”}, two noise levels {0.0%, 2%} and three fragment lengths {650 bp, 300 bp, 150 bp}.

### Deep learning model for taxonomic classification

---CNN model

The convolutional neural network (CNN) classification model employed a typical convolutional architecture used in multiple studies (Jiang et al. 2023; Zheng et al. 2019; Busai et al. 2019), consisting of convolutional blocks for feature extraction and subsequent fully connected (FC) classification layers. The convolutional part has three consecutive convolutional blocks with each sequentially consisting of the 1D convolution, batch normalization, 1D max pooling, rectified linear unit (ReLU, 𝑅𝑒𝐿𝑈(𝑥) = 𝑥 if 𝑥 > 0 otherwise 0), and dropout (rate = 0.15 for CNN layers and 0.25 for FC layers) layers. There were 64, 128, and 128 channels in the first, second, and third convolutional blocks, respectively. Hyperparameters, including the number of channels and dropout rates, were determined using cross-validation runs on a partial dataset.

An input DNA sequence with length L was encoded in an L × 4 matrix whose rows were four-dimensional one-hot vectors. For instance, a base letter “A” was represented as a row [1, 0, 0, 0], and “T” as [0, 1, 0, 0]. Noncanonical base letters (N, R, Y, etc.) were represented as [0, 0, 0, 0]. For each convolution process, the length was halved by one-dimensional (1D) max pooling with a size of two, resulting in an L/8 ×128-dimensional output. One-dimensional global average pooling was then applied to the outputs to obtain a 128-dimensional feature vector. Subsequently, classification was performed with three FC layers to classify the input sequences into known taxa. The softmax function was applied to the final output of the FC layer to obtain prediction probabilities. Details of the neural network architecture are presented in Supplementary figure S1.

Throughout the performance evaluation process, the models were trained using the Adam algorithm with a cross-entropy loss. Default hyperparameter settings provided by Keras were used (batch size=16, learning rate=0.001). The convergence of loss was visually assessed.

---Deep learning methods for OOD detection

In addition to the taxonomic classification model described above, we implemented deep learning methods for out-of-distribution (OOD) detection. OOD detection is the task of separating samples into two categories: *IN-DISTRIBUTION,* hereafter, *ID*, which includes samples from classes present in the training data, and *OUT-OF-DISTRIBUTION* or *OOD*, which includes samples from classes NOT present in the training data (Zhang et al. 2024). The accepted ID samples were subsequently classified into ID classes. We employed three methods based on the prediction uncertainty scores. We selected these methods based on their reported performances (Zhang et al. 2023) and implementation complexities. Methods designed to work without explicit OOD sample exposure during the training phase were selected, because OOD exposure is not feasible for real barcoding applications. We also excluded methods that required complex optimization of hyperparameters.

Three OOD scores were calculated from the output obtained from intermediate FC layers. When the output of the penultimate FC layers was *g(x)*, the following transformation to *g(x)* was applied in the final FC layer:

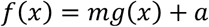

Here, *f(x)* is the output of the final FC layer, which is a vector of length equal to the number of classes; *m* is a weight matrix; and *a* is an offset vector. Each of these parameters were optimized in the training process. These intermediate outputs, *f(x)* and *g(x)*, contain useful information for discriminating OODs from ID samples (Supplementary figure S2).

--Maximum softmax probability

The maximum softmax probability (MSP) is commonly used as the prediction probability for neural network classification. The MSP score is defined as a function of the processes of exponentiation and scaling of *f(x)*, the output of the final FC layer, and its maximum value:

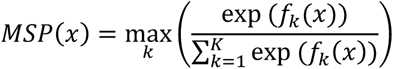

Here, 𝑓_𝑘_(𝑥) is the k-th component of the vector *f(x)*. The kth class that yields the MSP (i.e., argmax) is a predicted assignment of sample *x*. Importantly, this predicted class is chosen only from the classes present in the training dataset, regardless of whether the sample is of the OOD type. Hence, OOD detection is required to avoid the erroneous assignment of an OOD sample to a known class. Hendrycks and Gimpel (2016) proposed MSP as a metric for prediction uncertainty and showed that MSP scores of OOD samples were consistently lower than those of the ID sample, and a cutoff by a threshold of prediction probability helped to successfully detect OOD samples.

--Energy score

Liu et al. (2020) introduced the “energy score” of a neural network model for OOD detection. The log energy score of a neural network is defined as

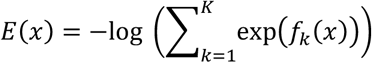

The log energy score is the logarithm of the softmax denominator in *MSP(x)*. The energy score is interpreted as the relative log-likelihood score of a model given a sample *x*, *Pr(x|model)*. The energy score of OOD samples was consistently lower than that of ID samples because the likelihood of obtaining such samples is less for models trained only with ID samples. Liu et al. (2020) reported that the threshold of sample energy values outperformed the softmax probability for multiple OOD detection tasks.

--Mahalanobis distance

Lee et al. (2018) developed a distance-based OOD detection method. The Mahalanobis distance of a sample from a class center is defined as

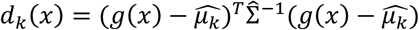

where 𝜇_𝑘_ is the k-th class center value, and Σ is a variance-covariance matrix of *g(x)*. Mahalanobis distance measures the distance from the k-th class center, assuming that the distribution of *g(x)* follows a multivariate normal distribution with a mean μk and a single variance-covariance matrix, Σ^^−1^, which are empirically estimated from a distribution of *g(x)* in a training data set. Lee et al. (2018) proposed the following negative Mahalanobis distance to the closest distribution center as an uncertainty metric for OOD detection:

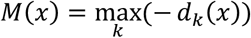

--Majority voting for OOD detection

In addition to independent OOD detection procedures with the above metrics, we devised a process for OOD detection with majority voting for the above three detectors. With this approach, a sample was treated as an OOD sample if two of the three methods “vote” for the presence of OOD.

All deep-learning models were implemented in Python using the Keras library. The code is available at https://github.com/tfujisawa/barcoding_cnn.

### Model training and performance test

The CNN models were trained with 70% of the ID data and their baseline prediction accuracy, the proportion of correct identifications to the total identification trials of the test samples, was measured. We then calculated the OOD scores (softmax probability, energy score, and Mahalanobis distance) for all ID test samples and obtained class thresholds by accounting for the 95% quantiles of all classes (Supplementary figure S3). After setting the thresholds, the model was exposed to OOD samples, and their scores were calculated. Samples with more extreme values than class-wise threshold values were classified as OODs. The proportion of OOD samples falsely classified as ID samples was measured as the false negative rate at a 95% threshold (FNR@95%). The training and evaluation processes were repeated 20 times for each dataset. The effects of fragment length, dataset size, noise level, and methods to determine the identification performance were assessed using multivariate linear regression. To assess the difficulty of the classification tasks, we calculated the proportion of misidentifications with zero genetic distances, i.e., the proportion of cases in which heterospecific specimens had identical sequences that led to misidentification of ID samples. The proportion of OOD specimens with zero genetic distances from any ID sample was also calculated. These zero-genetic-distance proportions determine the upper limits of classification accuracy and OOD detection (i.e., “perfect classifier”. Ziemski et al. 2021).

### Classification and OOD detection methods with distance

Conventional classification methods based on sequence distances were used as performance baselines. We measured the pairwise K2P genetic distance for the aligned sequences and the BLAST percentage similarity for the unaligned matrices (Altschul et al. 1990). Distance-based classification was then performed using the 1-nearest neighbor criterion (1NN), where a new sample was assigned to the taxon of a sample with the smallest distance from it. Although 1NN based on the K2P or BLAST distance is the simplest distance-based classification algorithm, it is still widely used and often outperforms more sophisticated algorithms (Leray et al. 2022; Hleap et al. 2021). For OOD detection tasks, we calculated the minimum distances from samples within their own class/species and set class OOD thresholds by taking the 95% quantiles of their minimum distances. When the distance between a test OOD sample and its nearest neighbor is greater than the class OOD threshold, the sample is classified as an OOD sample. The above procedure is similar to the “best close match” procedure proposed in Meier et al. (2006) although the quantile calculation process is different.

### Gradient-based attribution

We visualized the region responsible for classification decisions using a one-dimensional gradient-based class activation map (GradCAM, Selvaraju et al. 2016). GradCAM localizes the region of importance by measuring the effects of CNN features on the classification probabilities. Specifically, the GradCAM score on window *w,* is defined as

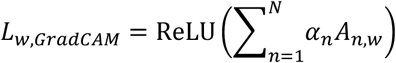

𝐿_𝑤,𝐺𝑟𝑎𝑑𝐶𝐴𝑀_ is a weighted average of *N* CNN features, 𝐴_𝑛,𝑤_, calculated on the window *w* with the weight 𝛼_𝑛_. The weight is a feature importance, measured as an averaged partial derivative of MSP(x) with respect to 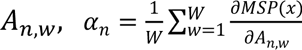, where *W* is the total number of windows on a sequence. An interpretation of importance is that when a unit change in a CNN feature (𝐴_𝑛,𝑤_) results in a significant change in the prediction probability (MSP(x)), 𝐴_𝑛,𝑤_ is considered to be important in the prediction process. We also implemented an activation map of the energy score to visualize the region responsible for OOD detection decisions by replacing the gradient of the MSP(x) in the weight calculation with the gradient of the energy score.

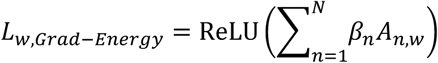

Here, the weight 𝛽_n_ is defined as 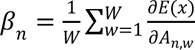. In this case, the effect of the CNN features on the energy score was measured. In the current study, the window size was set to 8 bp, resulting in 85 windows in a 680 bp fragment. We compared the GradCAM and Grad-Energy scores with genetic variations measured in 16 bp windows.

### Regression by population genetic metrics

A set of population genetic metrics was calculated for each genus dataset to identify the determinants of the identification performance. Genetic distance-related metrics, including average within-species and average and minimum between-species genetic distances, were calculated from alignments. Dataset completeness was measured by the average number of samples per species, the total number of species, and “taxonomic completeness,” defined by the number of ID species divided by the total number of all species. To identify factors affecting model performance, multivariate regression modeling was conducted using the above metrics as explanatory variables, and identification accuracy and false negative rate as responses. Least absolute shrinkage and selection operator (LASSO) procedures were used to select important explanatory variables.

## RESULTS

### BOLD dataset profiles

We collected 34,408 COI sequences from 20 genera in the BOLD database. Of the 13,078 examined genera in the four target orders, only 82 (0.6%) met the sample size criteria. The number of in-distribution (ID) and out-of-distribution (OOD) samples were 28,422 and 5,986, respectively. The number of species within the selected genera ranged from 15 to 68, and the average number of samples per species was 45. The number of OOD samples per genus ranged from 23 to 910, and the proportion of OOD samples to total samples was 0.17.

### Accuracy of identification and OOD detection

Both deep learning and distance-based methods were highly accurate in ID identification tasks, especially when trained with sufficiently large datasets. For whole 650 bp fragments, the average baseline prediction accuracy of the CNN model was 0.97, and the two conventional methods were as accurate as the CNN model (0.971 for k2p distance, 0.973 for BLAST; Fig. 2 and Table. 1). The accuracy decreased with reduced fragment sizes for all methods, and the CNN slightly outperformed the conventional methods when the fragment length was 150 bp (0.960 for CNN, 0.945 for k2p distance, and 0.946 for BLAST). Multiple linear regression analyses showed that shorter fragments were significantly associated with lower accuracy, and the CNN model exhibited slightly higher accuracy, but the difference was not significant. When the training datasets were smaller, the CNN performance decreased, whereas the distance methods were less affected. The introduction of 2% noise to the sequence reduced the identification performance; however, the reduction in accuracy was within 2% under most conditions (average accuracy decrease = 0.011, p<<0.001).

**Figure 1.**
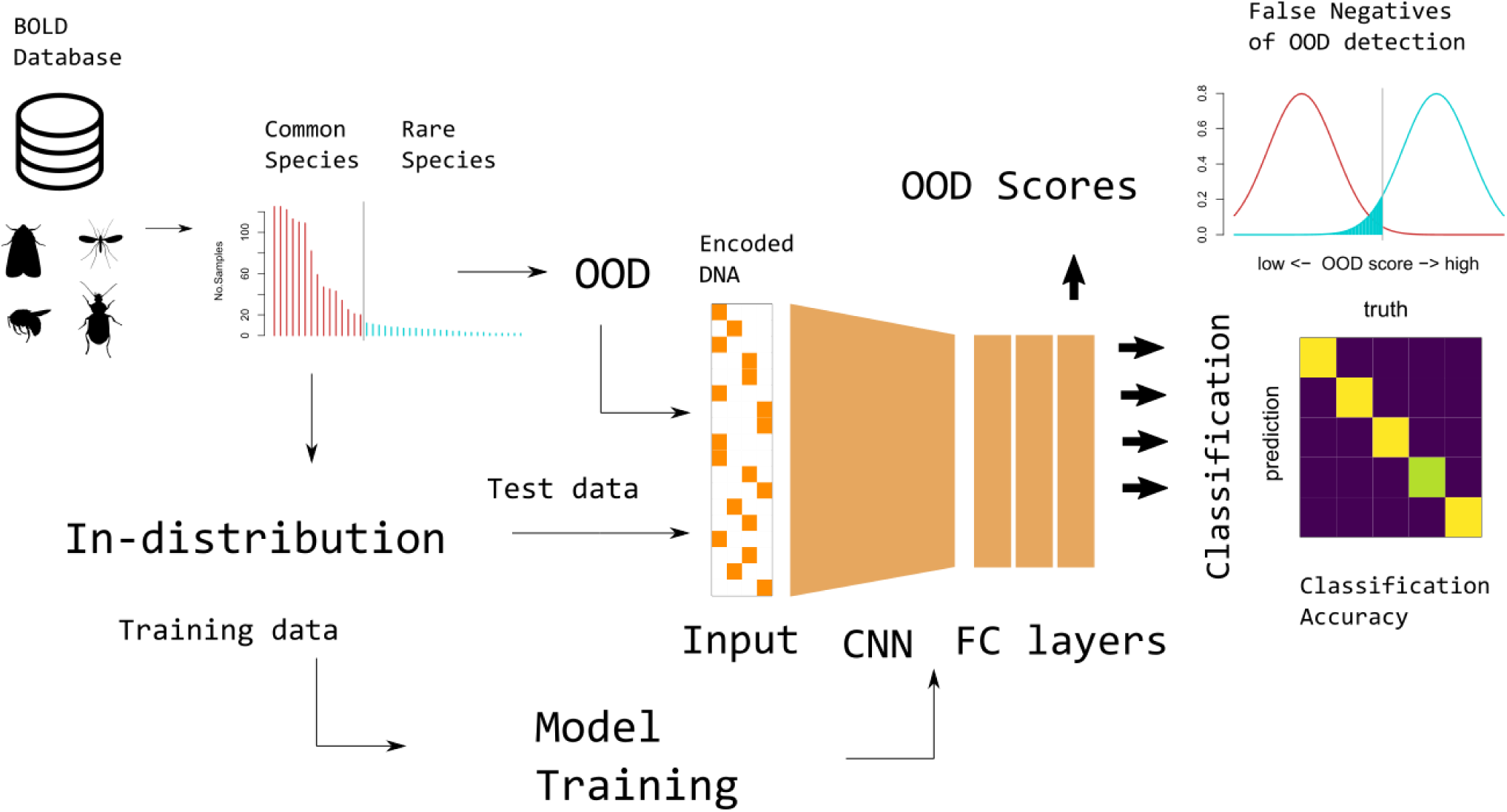
A schematic diagram of the classification model, data acquisition and analysis procedures.

**Figure 2.**
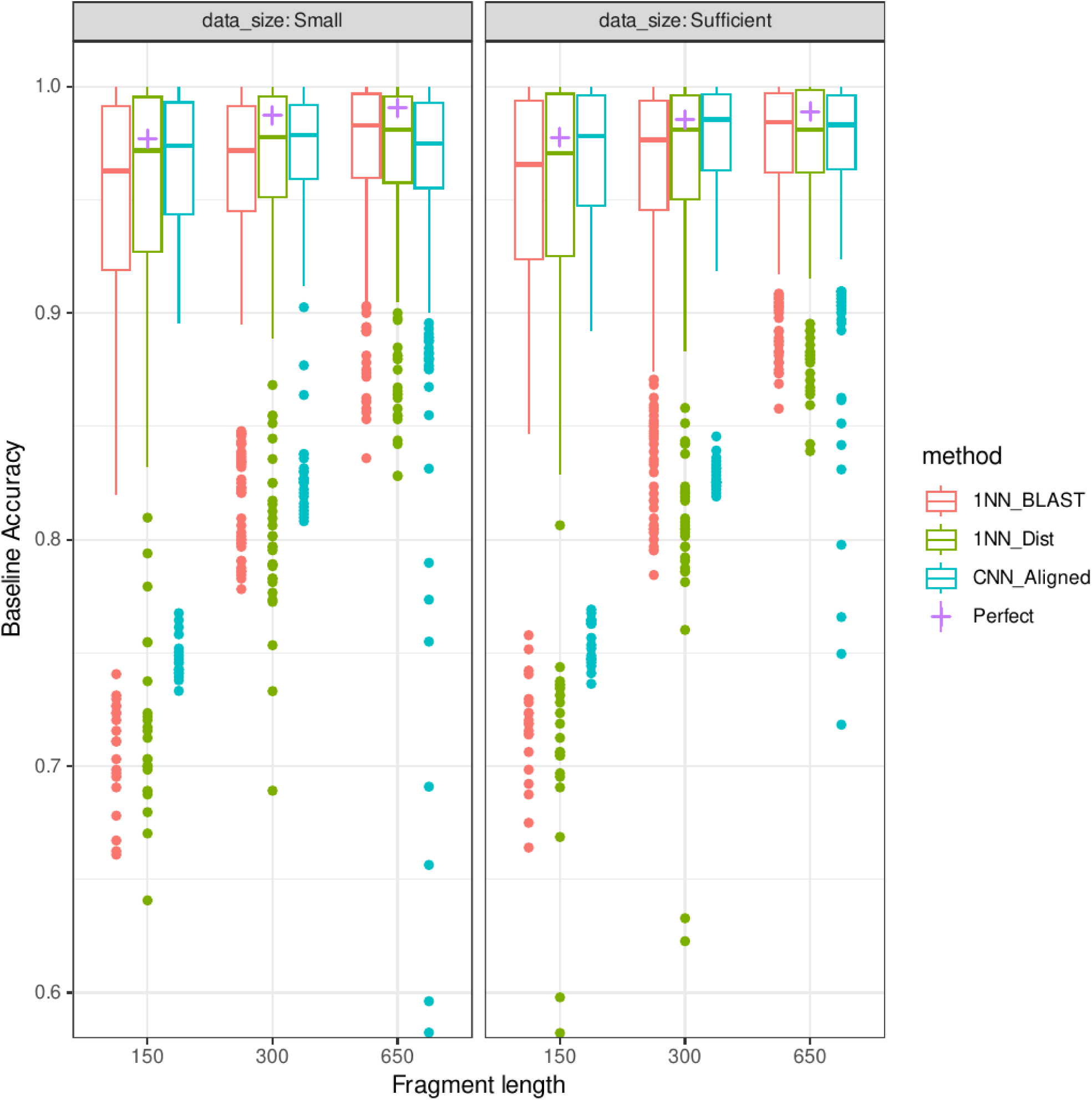
Effects of fragment lengths and database sizes on baseline prediction accuracy in the noiseless dataset.

The performance of OOD detection tasks generally exhibited patterns similar to those of identification tasks. The CNN model underperformed conventional methods with long fragments (FNR@95% 0.128 for CNN, 0.103 for k2p distance, and 0.101 for BLAST for 650 bp fragments; Figure 3 and Table 2) but outperformed with shorter fragments. However, the effect of reduced fragment size was more pronounced (FNR@95%: 0.156 for CNN, 0.176 for k2p distance, and 0.170 for BLAST for 150bp fragments). There was no significant difference in FNR between the detection methods. Among the deep learning methods for OOD detection, the performances of the energy score and Mahalanobis distance were closely matched, and these methods significantly outperformed MSP (Supplementary figure S4 and Table S2). Consensus across the three methods generally resulted in better performance, but the improvement was not significant, and the best-performing methods depended on the datasets. The proportion of OOD samples with zero distance from ID samples was 0.061 for 650 bp and reached 0.161 for 150 bp. The performance of the OOD detection method was close to these optimal values, although the gaps were greater for longer fragments (FNR_perfect_=0.061 vs. FNR_CNN_=0.128 for 650 bp and 0.156 vs. 0.161 for 150 bp).

**Figure 3.**
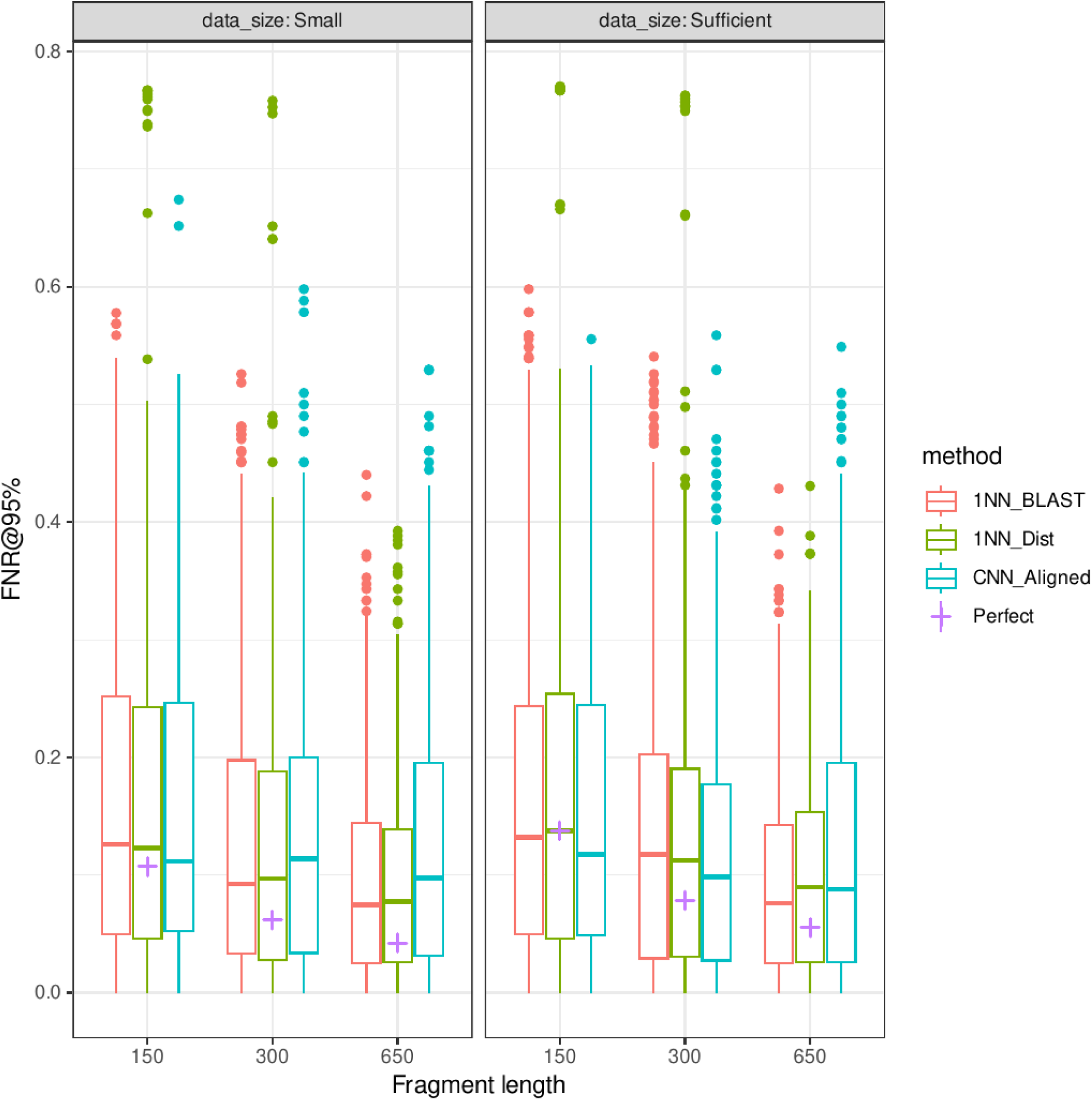
Effects of fragment lengths and database sizes on false negative rates of OOD detection in the noiseless dataset.

**Table 1.** Baseline prediction accuracy of three identification algorithms and the perfect classifier under different database sizes, noise levels, and fragment lengths. The best performing method in a set of parameters is indicated by values in boldface.

|  | Database Size | Sufficient |  |  |  | Small |  |  |  |
| --- | --- | --- | --- | --- | --- | --- | --- | --- | --- |
|  | Method | CNN | Distance | BLAST | Perfect | CNN | Distance | BLAST | Perfect |
| Noise Level | Fragment Length |  |  |  |  |  |  |  |  |
| 0 | 650 | 0.97 | 0.972 | **0.973** | 0.98 | 0.95 | 0.968 | **0.971** | 0.982 |
|  | 300 | **0.967** | 0.957 | 0.958 | 0.963 | **0.962** | 0.959 | 0.955 | 0.968 |
|  | 150 | **0.96** | 0.945 | 0.946 | 0.951 | **0.958** | 0.945 | 0.942 | 0.952 |
| 0.02 | 650 | 0.953 | 0.965 | **0.968** |  | 0.929 | 0.966 | **0.967** |  |
|  | 300 | **0.963** | 0.937 | 0.953 |  | **0.956** | 0.944 | 0.95 |  |
|  | 150 | **0.949** | 0.923 | 0.933 |  | **0.945** | 0.93 | 0.931 |  |

**Table 2.** False negative rates at a 95% threshold for three OOD detection methods and a perfect classifier under various parameter settings. A lower FNR indicates better performance. the best performing method in a set of parameters is indicated by values in boldface. For the CNN model, the results of majority voting (MV) with multiple methods are shown.

|  | Database Size | Sufficient |  |  |  | Small |  |  |  |
| --- | --- | --- | --- | --- | --- | --- | --- | --- | --- |
|  | Method | CNN(MV) | Distance | BLAST | Perfect | CNN(MV) | Distance | BLAST | Perfect |
| Noise Level | Fragment Length |  |  |  |  |  |  |  |  |
| 0 | 650 | 0.128 | 0.105 | **0.101** | 0.061 | 0.13 | **0.099** | 0.101 | 0.046 |
|  | 300 | **0.126** | 0.14 | 0.142 | 0.116 | 0.137 | **0.125** | 0.133 | 0.09 |
|  | 150 | **0.156** | 0.176 | 0.17 | 0.161 | **0.156** | 0.165 | 0.162 | 0.132 |
| 0.02 | 650 | 0.171 | 0.147 | **0.142** |  | 0.175 | **0.136** | **0.136** |  |
|  | 300 | **0.186** | 0.205 | 0.198 |  | 0.193 | **0.181** | 0.183 |  |
|  | 150 | **0.242** | 0.272 | 0.27 |  | **0.243** | 0.244 | 0.256 |  |

Sequences with noise significantly compromised the OOD detection performance for each method (Average FNR increase=0.064, p<<0.001). In reduced-size datasets, FNRs of the distance-based methods were slightly improved. The average *decrease* in the FNRs for small datasets over sufficient datasets was 0.0079, and linear regression analysis showed that the effect was significant (p=0.0026). These counterintuitive results are attributable to the reduced number of identical sequences shared between OOD samples and their nearest ID counterparts.

### Regression modeling

The results of the multiple regression analysis with LASSO variable selection are summarized in Table 3. Regression modeling showed that identification accuracy was positively correlated with the number of samples per species and the minimum between-species distance and negatively correlated with the number of classes and taxonomic completeness. FNR@95% was negatively correlated with the minimum between-species distance and positively correlated with the average within-species distance. A minor negative effect of the number of classes was also observed, while other variables were excluded. In addition, the identification accuracy and FNR were significantly correlated (Pearson’s r = -0.48, Figure 4), indicating that when the model correctly identified the ID classes, its OOD detection ability was accurate.

**Figure 4.**
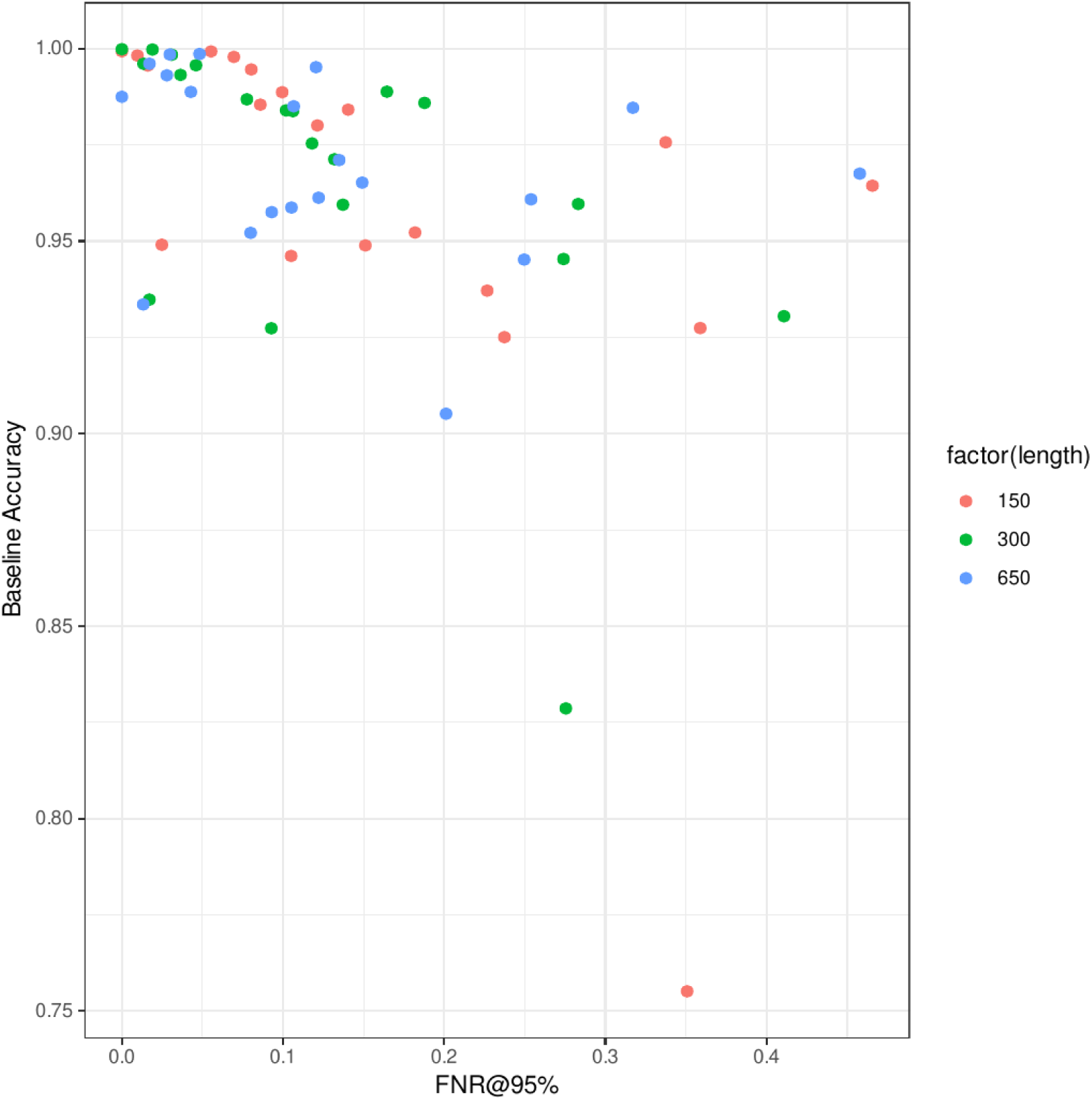
Relationship between the accuracy of the CNN classifier and the false negative rate of the CNN OOD detector.

**Table 3.** Regression coefficients estimated by multivariate regression modeling for baseline accuracy and false negative rate. Coefficients dropped by the LASSO variable selection are indicated by “.” signs. *Dw*: Within-species genetic distance. *Dbt* : Between-species genetic distance

|  | Explanatory | variables |  |  |  |  |
| --- | --- | --- | --- | --- | --- | --- |
| Response | No. classes | Average *Dw* | Average *Dbt* | Minimum *Dbt* | completeness | No. samples per species |
| Baseline Accuracy | -1.35e-05 | . | . | 0.040 | -0.051 | 6.39e-04 |
| False negative rate | -0.0007 | 1.031 | . | -1.16 | . | . |

### Gradient-based attribution

The sequence regions important for classification localized by GradCAM largely corresponded to regions with high genetic variation in the alignment. The genetic variations in 16 bp windows were strongly correlated with average GradCAM scores on the same windows (Pearson’s *r* = 0.41-0.76), and peaks were often aligned with the highest genetic variations.

This trend was consistently observed across fragment lengths (Figure 5). In contrast, a weaker correspondence was observed between the regions of importance in energy-based OOD detection and regions with high genetic variation (Figure 6), and the grad-energy score was less correlated with genetic variation (Pearson’s *r* = 0.06 - 0.45) that were always lower than those of GradCAM.

**Figure 5.**
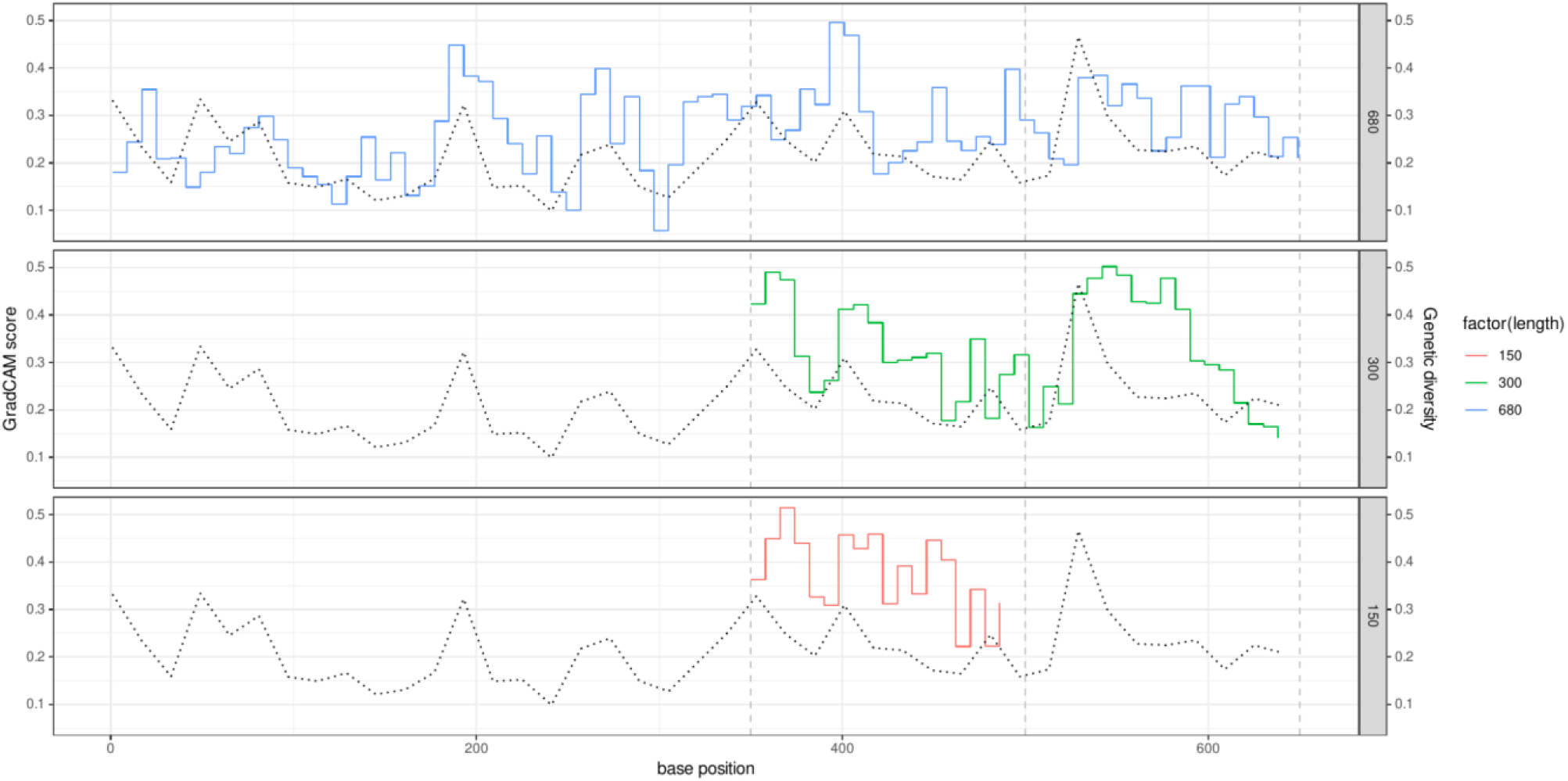
Spatial distribution of the average GradCAM score for in-distribution samples of the *Cryptocephalus* (leaf beetle) dataset. Solid step lines represent the GradCAM score for the 8-bp windows, and black dotted lines represent genetic variations in the 16-bp windows.

**Figure 6.**
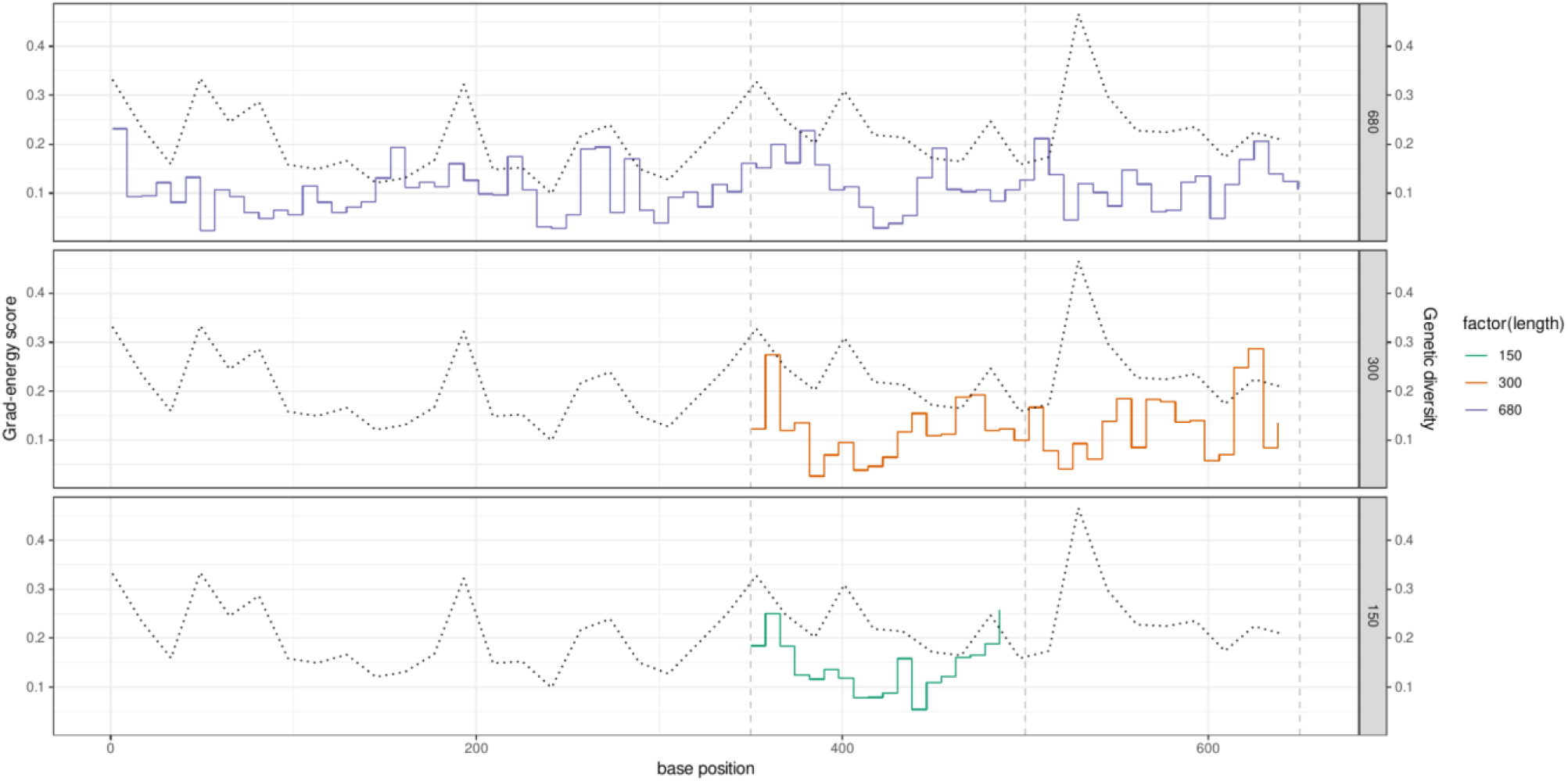
Spatial distribution of the average grad-energy scores for OOD samples from the *Cryptocephalus* data set. Solid step lines represent the grad-energy score for 8-bp windows, and black dotted lines represent genetic variations in 16-bp windows.

## DISCUSSION

Under the conditions considered in this study, sequence-based identification methods were highly accurate and robust against sequence noise when sufficient samples were available. The models slightly underperformed with short fragment lengths or smaller training datasets, as reported in previous studies (Porter & Hajibabaei 2018), but the best-performing models retained ∼95% accuracy. Regardless of minor performance differences, DNA barcoding identification methods appear to have already been optimized, and their performance is very close to that of the ideal classifier. The reduction in accuracy was largely due to the reduced number of diagnostic variations, as reported by Ziemski et al. (2021).

By contrast, sequence-based out-of-distribution (OOD) detection was a more refractory task, with higher error rates for short and noisy fragments. In addition, the performance was more counterintuitively dependent on the database size (e.g., improved FNR@95% with *smaller* databases). More importantly, the accuracy of OOD detection was more strongly limited by samples that were undiagnosable by sequencing alone, comprising ∼16% of OOD samples under some extreme conditions. Although there is room for improvement in OOD detection using long fragments and smaller databases, a strong limiting factor is the lack of diagnostic characters for short fragments.

The risk of overlooking OOD samples has been recognized but considered difficult to quantify (Virgilio et al. 2010). This study provides a coarse estimate of these risks. Assuming that the current proportion (17%) of bulk specimens is of the OOD type, up to ∼3% of the total specimens may be misidentified as referenced species, with errors increasing when targets containing more unknown samples or noisy sequences are used. Under such conditions, sequence-based surveys may significantly underestimate unknown biodiversity even with the best identification methods. Because most insect species have not been barcoded and are highly likely to have variations that will be missed during fragment truncation, it is prudent to use as long fragments as possible to minimize the risk of overlooking them. Because the performance of OOD detection was correlated with within-species distance and minimum interspecific distance, these metrics may be used to help determine the appropriate marker length. In addition, the correlation between ID classification and OOD detection performance can be used to assess potential risks and improve detection performance during the training process (Vaze et al. 2021).

In this study, we compared distance-based methods with deep learning. Although their overall performances were similar, the general tendency was that short fragments favored the CNN model, and longer fragments favored the distance methods. Deep learning classification performed poorly with small database sizes and long fragment lengths. This performance reduction may reflect difficulties in optimizing highly parameter-rich models (∼110k parameters). Our deep learning OOD detection methods exhibited a performance comparable to that of the distance methods. Therefore, they may be used to mitigate the reported performance reduction of deep learning models due to incomplete training databases (Romeijn et al. 2024).

Deep learning models also provide useful information for interpreting results. Gradient-based explanations of CNN classification showed that highly variable regions in the alignment were informative for the classification tasks of in-distribution (ID) samples but not necessarily for OOD detection tasks. This result may reflect the different natures of the two tasks. For example, in an extreme case, a site might be completely invariable among ID samples, while an OOD species harbors a diagnosable difference at the same site. Under these conditions, the site is uninformative for classification but highly informative for OOD detection. Such site importance score may be useful for distinguishing favorable sites for different tasks and for selecting informative markers.

We applied only a limited number of OOD detection methods in this study, and methodological improvements may exist. For example, training models with ID and synthetic OOD samples may potentially improve detection performance. Nevertheless, as taxonomic coverage and within-species sample sizes improve, identification success will depend more on sequence variation than on identification procedures. Except for using longer fragments, a straightforward improvement is sequencing multilocus markers to increase the available diagnosable variations. Although sequencing short multilocus markers can be cost-effective (Wang et al. 2023), they may result in incongruent species compositions even in a single community because of different PCR affinities. An alternative approach to OOD tolerant identification is to use additional information, such as geographic locations, environmental information, and morphological features. For example, fine-grained geographic information can not only be used to identify species, but also to detect possible OOD samples because insect communities can have extremely high geographic turnover (Srivathsan et al. 2023; Arribas et al. 2021). When closely related species occupy different niches, environmental niche modeling can provide additional information for identification (Yang et al. 2024). A similar approach may be used for OOD detection. High-throughput imaging is another potentially useful tool to supplement metabarcoding (Fujisawa et al. 2023; Wührl et al. 2022) and has been successfully used to verify the metabarcoding results (Panel et al. 2025). Machine learning algorithms may help integrate multiple information sources because designing “multimodal” models combining different types of data is easier than using conventional statistical methods.

In summary, sequence-based identification with DNA barcoding is highly accurate, owing to collective efforts for performance improvement. However, incomplete databases and the presence of OOD samples still pose methodological challenges, and a careful experimental design is required to avoid overlooking these unknowns. In the future, machine learning models should integrate multiple sources of information for more robust and unknown-proof identification. The rapid accumulation of DNA sequence databases and additional ecological information, along with advanced machine learning algorithms, may enable the deployment of integrated biodiversity monitoring systems.

## Acknowledgements

This study was supported by JSPS KAKENHI (Grand No. 20K06824).

## Supplementary figures

**Supplementary figure S1.**
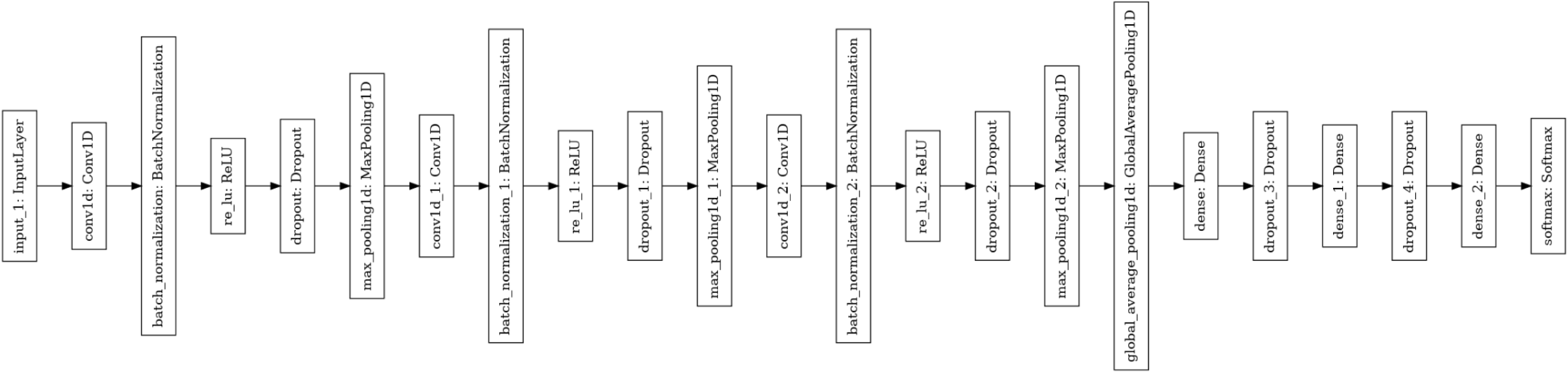
A diagram showing the detailed architecture of the CNN model.

**Supplementary figure S2.**
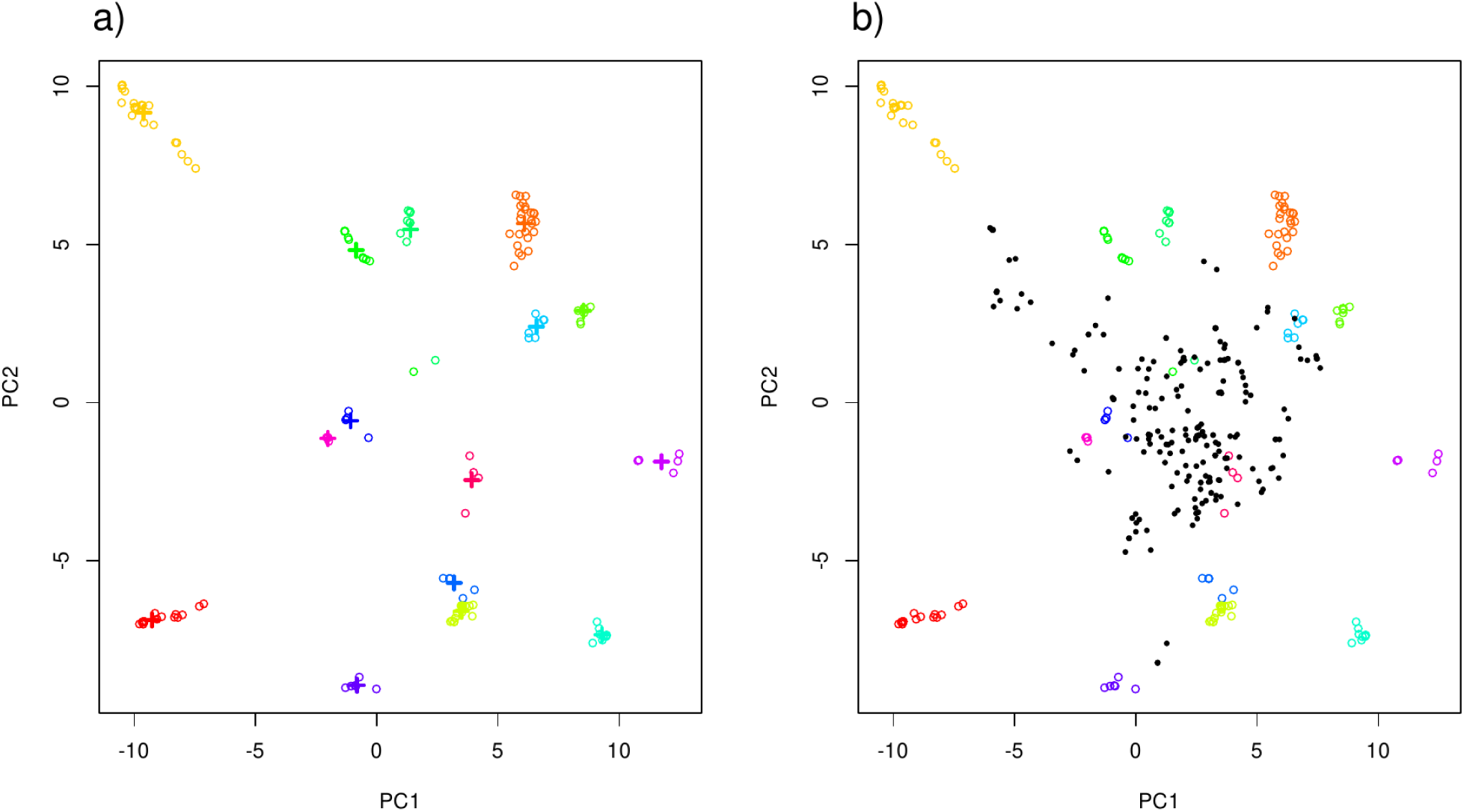
Exemplary distribution of *g(x)*, showing the outputs of the penultimate FC layer. PCA was applied to reduce the dimensionality for visualization. a) Dots in colors represent in-distribution (ID) samples of different species, while crosses in corresponding colors are class centers; μk. b) the same plots with OOD samples are shown in black dots. ID samples were frequently clustered in linearly separable groups in the intermediate output space, while OOD samples were placed between such groups. Hence, distances from class centers to samples can be used to measure the OOD status of samples.

**Supplementary figure S3.**
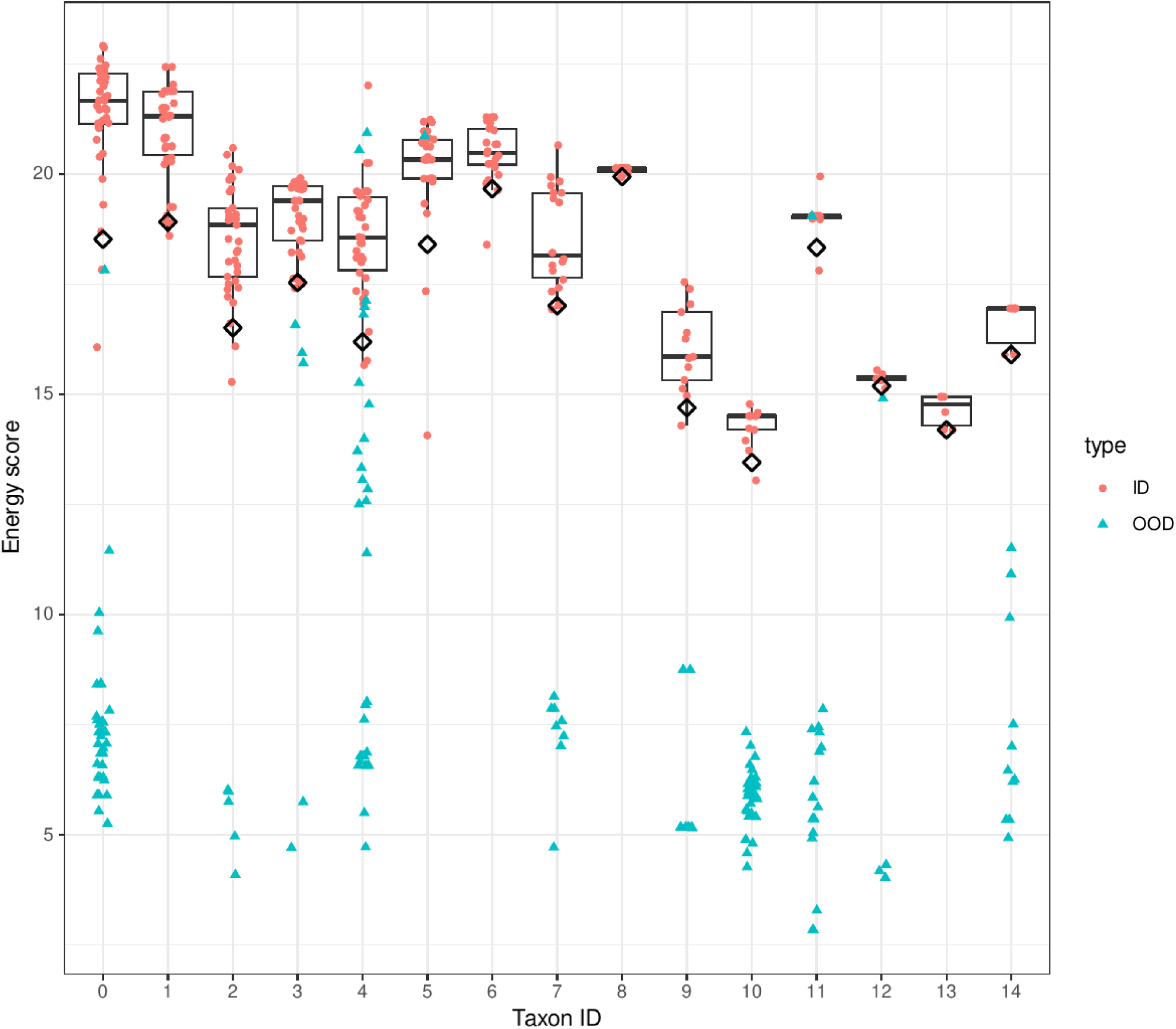
Distribution of energy scores for ID and OOD samples from the Drosophila dataset. Taxon IDs of the OOD samples were assigned based on the predictions of the CNN classifier. Open squares indicate the 95% quantiles of the energy scores of ID samples. Samples with lower energy scores with these thresholds were detected as OODs. OOD samples missed by these procedures, such as those with high energy scores in Taxon ID 4, were considered false negatives.

**Supplementary figure S4.**
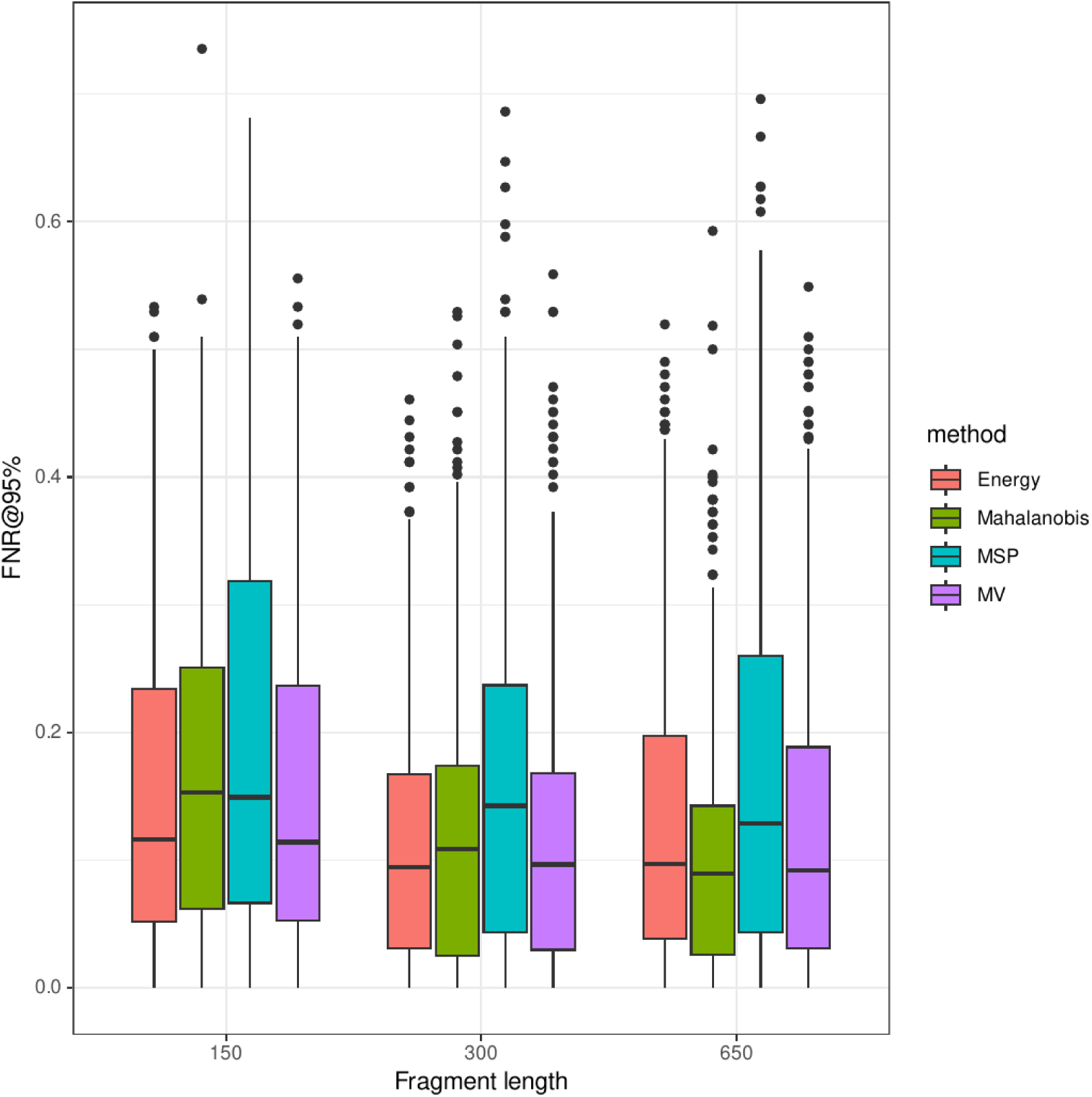
False negative rates (FNR@95%) of four OOD detection methods and their relationships with fragment lengths. Results on the noiseless sufficient-size dataset are shown. MSP: Maximum Softmax Probability, MV: Majority Voting.

**Supplementary table 1.** Dataset summary.

| Genus | Dataset Code | Order | Common Name | No.ID.samples | No.ID-species | No.OOD samples |
| --- | --- | --- | --- | --- | --- | --- |
| *Drosophila* | dro15 | Diptera | fruit fly | 1080 | 15 | 162 |
| *Megaselia* | meg42 | Diptera | scuttle fly | 2296 | 42 | 348 |
| *Aedes* | aed19 | Diptera | tiger mosquito | 1169 | 19 | 102 |
| *Atheta* | ath18 | Coleoptera | rove beetle | 707 | 18 | 228 |
| *Cryptocephalus* | cry22 | Coleoptera | leaf beetle | 828 | 22 | 304 |
| *Pterostichus* | pte45 | Coleoptera | ground beetle | 2383 | 44 | 169 |
| *Dolerus* | dol24 | Hymenoptera | sawfly | 907 | 24 | 135 |
| *Megachile* | mch15 | Hymenoptera | leafcutter bee | 649 | 15 | 240 |
| *Lassioglossum* | las68 | Hymenoptera | sweat bee | 2960 | 68 | 910 |
| *Euxoa* | eux40 | Lepidoptera | owlet moth | 1623 | 40 | 778 |
| *Phyllonorycter* | phy53 | Lepidoptera | leaf mining moth | 2086 | 53 | 439 |
| *Acleris* | acl32 | Lepidoptera | leaf roller moth | 1645 | 32 | 138 |
| *Culicoides* | cul26 | Diptera | biting midge | 992 | 26 | 248 |
| *Amara* | ama25 | Coleoptera | sun beetle | 1142 | 25 | 260 |
| *Catocala* | cat62 | Lepidoptera | underwing moth | 2135 | 62 | 355 |
| *Andrena* | and28 | Hymenoptera | mining bee | 873 | 28 | 565 |
| *Bombus* | bom24 | Hymenoptera | bumble bee | 1187 | 24 | 167 |
| *Corynoptera* | cor19 | Diptera | fungus gnat | 1927 | 19 | 23 |
| *Bembidion* | bem39 | Coleoptera | ground beetle | 1053 | 39 | 226 |
| *Caloptilia* | cal17 | Lepidoptera | leaf mining moth | 780 | 17 | 189 |

**Supplementary table 2.** False negative rates at the 95% threshold for the four OOD detection methods of the deep learning model. The best performing methods are indicated in boldface. The results for a noiseless, sufficiently sized dataset are shown. MSP: Maximum Softmax Probability, MV: Majority Voting.

|  | Database | Sufficient |  |  |  |
| --- | --- | --- | --- | --- | --- |
|  | Method | MSP | Energy | Mahalanobis | MV |
| Noise level | Fragment length |  |  |  |  |
| 0.0 | 650 | 0.169 | 0.13 | **0.11** | 0.128 |
|  | 300 | 0.171 | **0.124** | **0.124** | 0.126 |
|  | 150 | 0.202 | 0.157 | 0.167 | **0.156** |

